# Controlling heterologous protein synthesis through a plant RNA ThermoSwitch

**DOI:** 10.1101/2024.10.07.616989

**Authors:** Filip Lastovka, Hadrien Peyret, Sherine Elizabeth Thomas, George P. Lomonossoff, Betty Y.-W. Chung

## Abstract

Plants have evolved sophisticated mechanisms to adapt to temperature fluctuations, including transcriptional, post-transcriptional, and post-translational processes. Recent discoveries highlight RNA ThermoSwitches, cis-acting elements in several plant mRNAs that regulate protein synthesis based on temperature changes. These mechanisms, first identified in Arabidopsis thaliana, offer a promising tool for biotechnology by enabling temperature-sensitive control of protein expression. This study demonstrates, for the first time, the feasibility the application of plant RNA ThermoSwitches in Agrobacterium-mediated transient expression systems, presenting a novel method for controlled gene expression in plants. This system is particularly advantageous due to its homogeneous nature and independence from chemical inducers or suppressors.

## Main text

Plants have evolved mechanisms to adapt to fluctuations in surrounding temperatures, to which they are constantly exposed. Many such mechanisms operating at transcriptional, post-transcriptional, and post-translational levels have been extensively investigated [1] [2]. Recent studies have revealed the existence of a direct translational control mechanism that operates *in cis* in several plant mRNAs. These RNA ThermoSwitches, first discovered in *Arabidopsis thaliana*, act as temperature sensors to directly regulate protein synthesis from the associated mRNAs [3] [4].

One of the best characterized plant RNA ThermoSwitches is an mRNA hairpin embedded in the 5′ UTR of the transcription factor gene *PIF7* (Phytochrome-Interacting Factor 7). The *PIF7* RNA ThermoSwitch is located 30 nt upstream of the translation start site (figure 1b) and directly modulates synthesis of the PIF7 protein in response to temperature. Below 22 °C the RNA ThermoSwitch is in a stable hairpin conformation that hinders the scanning pre-initiation complex (PIC) from reaching the translation initiation site. Upon temperature rise to 27–32 °C, the hairpin adopts a relaxed conformation enabling progression of PIC scanning until it reaches the initiation site for protein synthesis (figure 1a) [3] [4]. At the same time, the partially folded, yet relaxed ThermoSwitch temporarily obstructs the incoming PIC, enabling the preceding complex to initiate protein synthesis. The PIF7 protein subsequently drives the transcription of genes including the auxin biosynthesis gene cluster, thereby mediating day-time rhythmic growth and development of the plant in response to the temperature environment [3] [4].

**Figure 1.**
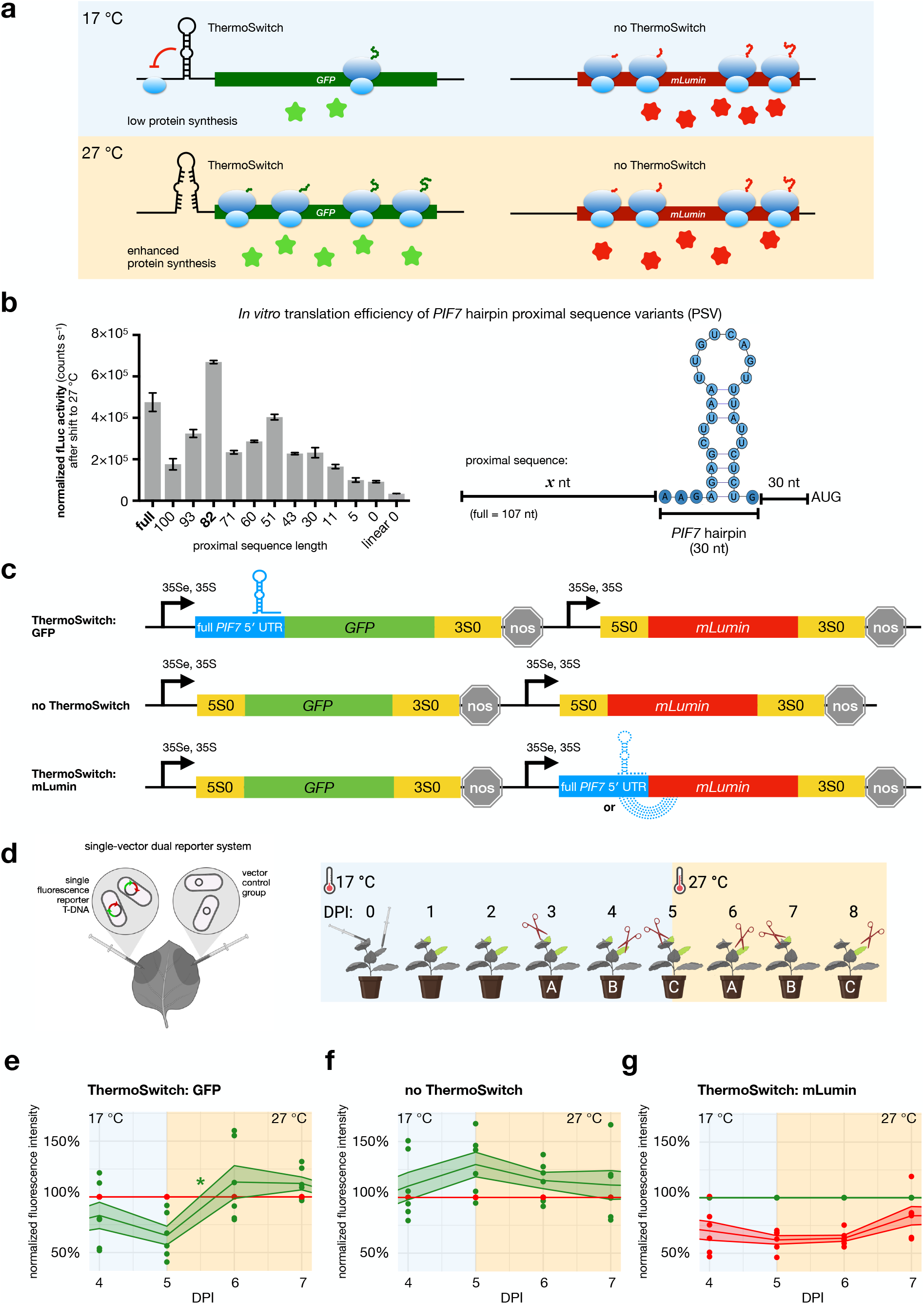
**a**, Regulation of protein synthesis by the ThermoSwitch before and after a temperature shift. **b**, *In vitro* translation efficiencies of PIF7 5′ UTR mRNA hairpin proximal sequence variants compared to the wild-type sequence after a 17-to-27 °C shift. Firefly luciferase activity normalized to corresponding mRNA levels is plotted on the *y*-axis. The error bars show the SEM. The RNA secondary structure was predicted with *Vienna RNAfold* [8] and visualized using *forna* [9]. **d**, Infiltration of *Agrobacterium* suspensions into *N. benthamiana* leaves and the time course experiment. The infiltrated leaves are highlighted in green. Scissors indicate leaf harvest, which rotated among three groups (A, B, C) of six plants. The subfigure was created with BioRender.com. **e**, Relative green fluorescence intensity (*PIF7* full ThermoSwitch-containing UTR) normalized to the red fluorescence intensity (5S0, without any ThermoSwitch). The asterisk represents a significant difference following the temperature shift: *p* = 0.0177. **f**, Relative green fluorescence intensity (5S0) normalized to the red fluorescence intensity (an identical 5S0). **g**, Relative red fluorescence intensity (PIF7 full, but alternative secondary structure) normalized to the green fluorescence intensity (5S0). The orange area represents a temperature of 27 °C. The ribbon shows the mean and the SEM. Differences between neighbouring time points without an asterisk were not statistically significant.

To further understand the effect of PIC scanning on the ThermoSwitch mechanism, we designed a *PIF7* 5′ UTR proximal sequence deletion series fused to the firefly luciferase coding sequence (fLuc; figure 1b). This enabled us to monitor translation efficiencies of *PIF7* proximal sequence variants (PSVs) compared with the full-length wild-type *PIF7* 5′ UTR in a wheat-germ *in vitro* translation experiment. With the exception of the 82-nt PSV, we observed an overall decrease in translation efficiency with decreasing 5′ UTR length in response to temperature shift from 17 °C to 27 °C. This reduction in protein synthesis with lower mRNA leader length could possibly be attributed to decreased preloading of scanning 40S subunits as noted previously in yeast and mammalian systems [5] [6]. Although further analysis will be needed to understand why the 82-nt PSV led to a 30% increase in translation efficiency from the full-length sequence, a secondary structure prediction indicates the presence of an additional weaker hairpin formation at the 5′ end of this mRNA. Interestingly, all deletion variants including a complete deletion of 5′ UTR proximal sequence of the *PIF7* hairpin resulted in enhanced translation at higher temperature in comparison with an unstructured RNA sequence of the same length and nucleotide content (linear 0; figure 1b). This indicates that although scanning by PIC is an important determinant in the control of *PIF7*-mediated translation initiation, the hairpin alone is *bona fide* ThermoSwitch that drives enhanced translation in response to increased temperature.

The ability to control protein expression levels by simply shifting the temperature at which plants are maintained is an attractive application in biotechnology, particularly in instances where high-level constitutive production of the target protein has undesirable effects. We have therefore investigated whether the plant RNA ThermoSwitch can be applied for temperature-sensitive control of gene expression in combination with Agrobacterium transient expression (figure 1a).

Fluorescent reporters were used to test the effect of a 5^′^ UTR ThermoSwitch on protein synthesis in *Nicotiana benthamiana* subjected to a 10 °C temperature shift. A dual reporter system was developed in the pHRE *Agrobacterium*-mediated expression vector to normalize the ThermoSwitch-dependent expression using the control reporter gene downstream of an artificially designed linear 5′ UTR (5S0) [7]. The two reporter genes were located in the same T-DNA region and possessed identical promoters (figure 1c). The capacity of the expression system to produce discrete fluorescence signals corresponding to both reporters that was measurable above background was verified in *Agrobacterium*-infiltrated plants grown in a glasshouse (figures S1a and b). Due to potential alternative base pairing of the ThermoSwitch with the *mLumin* coding region (figure S1c), *GFP* was chosen as the reporter of ThermoSwitch-dependent expression.

The dual reporter construct was introduced by *Agrobacterium* infiltration into leaves of five-week-old pre-flowering *N. benthamiana*. The plants were grown at 17 °C for five days prior to 27 °C shift, followed by harvesting at designated time points as illustrated on figure 1d for fluorescence quantification. Additionally, two controls were included: a negative control where both reporters were fused to 5S0 5′ UTR was included to further confirm ThermoSwitch-dependent translation and a construct where the reporters were switched (ThermoSwitch fused to *mLumin* instead of *GFP*) to confirm the inhibitory effect of potential base pairing between the ThermoSwitch and the *mLumin* coding region. As expected, significant increase in relative ThermoSwitch-controlled reporter signal was apparent 24 h post shift to 27 °C (figure 1e), while the relative fluorescence of the negative control remained constant despite higher temperature (figure 1f). The potential base pairing between the ThermoSwitch and *mLumin* RNA sequence significantly impaired the mLumin signal (figure 1g), highlighting the importance of careful assessment of potential RNA base pairing in designing expression systems.

In summary, for the first time, we have demonstrated the applicability of an RNA ThermoSwitch in *Agrobacterium*-mediated transient expression. The findings mark a step towards the development of an engineered plant thermosensitive expression system with potential application in molecular farming or in engineered crop plants to increase their yield or fitness [4]. A specific benefit of thermosensitive regulation is its homogeneous nature and its independence from chemical inducers or suppressors. Further work will focus on engineering ThermoSwitch sequences to achieve an augmented enhancing or suppressive effects through temperature-controlled translational induction.

## Authors’ contributions

B.Y.W.C and G.P.L. conceived the research. F.L., H.P., S.E.T., G.P.L., and B.Y.W.C designed experiments. S.E.T. performed all *in vitro* translation experiments. F.L performed all transient expression *in planta* experiments. F.L., H.P., S.E.T., G.P.L., and B.Y.W.C wrote the manuscript.

## Acknowledgements

We thank Dominykas Murza for contributing to the production of the reporter constructs and the Horticultural Services Department at the John Innes Centre for plant growth and Keith Saunders for day-to-day lab support.

This work was supported by BBSRC project grant BB/V006096/1 to B.Y.W.C and G.P.L.; A Medical Research Council Fellowship [MR/R021821/1] and BBSRC project grants [BB/X001261/1, BB/V017780/1] to B.Y.W.C. At JIC, the work was additionally supported by the BBSRC Institute Strategic Programme Grant “Harnessing Biosynthesis for Sustainable Food and Health” (BB/X01097X/1) and the John Innes Foundation.

## Supporting information

**Figure S1:** Supplementary figures

**Table S1:** *PIF7* full *GFP* and 5S0 *mLumin* fluorescence measurements

**Table S2:** 5S0 *GFP* and 5S0 *mLumin* fluorescence measurements

**Table S3:** 5S0 *GFP* and *PIF7* full *mLumin* fluorescence measurements

**Table S4:** 5′ UTR and primer sequences

## Appendix

Methods

## References

[1] J. J. Casal and S. Balasubramanian, “Thermomorphogenesis,” Annual review of plant biology, vol. 70, p. 321–346, 29 April 2019.

[2] S. Hayes, J. Schachtschabel, M. Mishkind, T. Munnik and S. A. Arisz, “Hot topic: Thermosensing in plants,” Plant, Cell & Environment, vol. 44, no. 7, p. 2018–2033, July 2021.

[3] B. Y. W. Chung, M. Balcerowicz, M. Di Antonio, K. E. Jaeger, F. Geng, K. Franaszek, P. Marriott, I. Brierley, A. E. Firth and P. A. Wigge, “An RNA thermoswitch regulates daytime growth in Arabidopsis,” Nature Plants, vol. 6, p. 522–532, 2020.

[4] S. E. Thomas, M. Balcerowicz and B. Y.-W. Chung, “RNA structure mediated thermoregulation: What can we learn from plants?,” Frontiers in Plant Science, vol. 13, 17 August 2022.

[5] M. Kozak, “Effects of long 5′ leader sequences on initiation by eukaryotic ribosomes in vitro,” Gene expression, vol. 1, no. 2, p. 117, 1991.

[6] J. J. van den Heuvel, R. J. Bergkamp, R. J. Planta and H. A. Raué, “Effect of deletions in the 5′-noncoding region on the translational efficiency of phosphoglycerate kinase mRNA in yeast,” Gene, vol. 79, no. 1, p. 83–95, 1989.

[7] H. Peyret, J. K. M. Brown and G. P. Lomonossoff, “Improving plant transient expression through the rational design of synthetic 5′ and 3′ untranslated regions,” Plant methods, vol. 15, p. 1–13, 2019.

[8] L. Ronny, S. H. Bernhart, C. Höner zu Siederdissen, H. Tafer, C. Flamm, P. F. Stadler and I. L. Hofacker, “ViennaRNA Package 2.0,” Algorithms for molecular biology, vol. 6, p. 1–14, 2011.

[9] P. Kerpedjiev, S. Hammer and I. L. Hofacker, “Forna (force-directed RNA): Simple and effective online RNA secondary structure diagrams,” Bioinformatics, vol. 31, no. 20, p. 3377–3379, 22 June 2015.

