## Supplementary Information for "Controlling heterologous protein synthesis through a plant RNA ThermoSwitch"

#### Methods

##### Design of the reporter system

The *Agrobacterium*-mediated transient expression plasmid pHRE [1] was used as a backbone for the reporter constructs. The *GFP* reporter gene originated from pHRE-GFP [1]. The red fluorescence gene used for the reporter system was *mLumin* [2] inserted into pHRE at the *Bsa*I restriction sites. *PIF7*-derived 5′ UTRs were inserted into the constructs at the *Bsm*BI restriction sites, replacing the original 5S0 5′ UTR, but retaining the Kozak sequence. To generate the dual reporter construct, the *GFP* reporter gene—which consists of the 35Se enhancer, 35S promoter (*Cauliflower mosaic virus*), the UTRs, the CDS, and the *nos* (nopaline synthase) terminator—was amplified by PCR (Phusion, Thermo Scientific) with primers that produced *Sac*I restriction sites at each terminus (table S4). This gene was then inserted into the *mLumin* reporter plasmid at the *Sac*I restriction site in the same orientation as *mLumin*. The reporter gene in the resulting construct possessed identical promoters, 3′ UTRs, and terminators.

A control *Agrobacterium* suspension was included in the experiments, which carried the empty vector and transferred a silencing suppressor and kanamycin resistance genes, but no fluorescence genes. This enabled confirmation that any autofluorescence from the tissue or the defence response to infiltration was not interpreted as the reporter signal.

##### Plant growth and infiltration

*Agrobacterium* carrying a reporter plasmid or the empty pHRE was cultured overnight from single colonies in 20–30 mL of LB supplemented with 50 μg $mL^{-1}$ of kanamycin and 50 μg $mL^{-1}$ of rifampicin in a 250 mL conical flask with a spring coil at 28 °C and 220 rpm. The overnight culture was pelleted by centrifugation in 50 mL tubes at a speed of 3.6 krcf and the supernatant was discarded. The pellet was resuspended and diluted in infiltration buffer to $OD_{600}$ 0.4. The infiltration buffer consisted of 10 mM 2-morpholin-4-ylethanesulfonic acid (MES) pH 5.6, 10 mM $MgCl_2$, and 100 μM acetosyringone. The suspensions were mixed in the required combinations and incubated at room temperature for more than 1 h before infiltration.

The *Agrobacterium* suspension was infiltrated using a needleless syringe into leaves of potted five-weeks-old pre-flowering *N. benthamiana* cultivar LAB. A small scratch was made on the epidermis of the adaxial surface of the leaf attached to the plant. The tip of the needleless syringe was pressed against the scratch and a finger was gently pressed on the opposite side of the leaf to ensure injection of the suspension into the leaf without causing excessive damage. As the suspension was injected, the infiltrated area was distinguishable as an expanding dark stain. The injection was continued until the infiltrated area no longer expanded or sufficient area was infiltrated. Further locations on the leaf were scratched and injected if a greater infiltrated area was required. The outlines of the infiltrated areas were marked with a marker.

Each leaf was infiltrated with the reporter suspension as well as the empty vector suspension in separate non-overlapping areas (figure 1d, left). Eighteen plants were grown in a growth cabinet (Sanyo, MLR-352H-PE) set to a constant temperature of 17 °C and a light cycle with an 8-h-long darkness and a 16-h-long maximal light intensity. On day five post infiltration (5 DPI) after the leaf harvest, the temperature was increased to 27 °C (figure 1d, right).

**Harvest and lysis of leaves**

The plants were assigned into three groups of six, with individuals of each group evenly distributed in the growth cabinet. Starting three days post infiltration (DPI), a single leaf of all plants in a single group was harvested by severing it at the petiole. At each time point, equal number of younger and older leaves of the two infiltrated leaves in each plant were harvested. The harvest rotated to another group the following day.
From each infiltrated area, three discs were extracted using a cork borer with a 12 mm diameter. The discs were placed into a shatterproof 2 mL centrifuge tube, snap-frozen in liquid nitrogen, and stored at –80 °C. The samples were further processed after the end of the time course. A ceramic bead and 270 µL of phosphate-buffered saline (PBS) with cOmplete EDTA-free protease inhibitor (Roche) were added to each tube. The discs were lysed using an Omni Bead Ruptor 24 homogenizer (settings: 30 s, single cycle, and maximal speed of 4 m $s^{-1}$). To obtain the soluble fraction, the lysate was centrifuged at 16 krcf in the cold room for 10 min, followed by transfer of 240 µL of the supernatant to a centrifuge tube and a second round of centrifugation.

**Fluorometry**

The clarified lysate was transferred to three wells of a 96-well opaque F-bottom black microplate, with 60 µL in each. The fluorescence was measured on a CLARIOstar Plus microplate reader (BMG LABTECH) using the top optic and 50 flashes per well. The excitation/emission wavelengths used were 470-15/515-20 nm to detect GFP and 585-15/630-20 nm to detect mLumin. To enable cross-plate comparisons, the gain was adjusted, separately for each experiment, at approximately 90% capacity using the most fluorescent well with lysate from 8 DPI (last day of the time course) and used for fluorescence measurement in all plates.

To calculate the relative fluorescence intensity values, fluorescence intensity (mean of the fluorometry measurements in three wells) of the ThermoSwitch reporter was divided by the fluorescence intensity of the reference reporter separately for each sample. The statistical significance of change in relative fluorescence intensity in the following time point was determined using Student's two-sample two-sided *t*-test. The normal distribution of the data was verified with the Shapiro–Wilk test and the homogeneity of variance was verified by Bartlett's test.

##### ***In vitro* translation experiments**

*In vitro* translation assays were performed using *PIF7*-derived 5′ UTRs fused to firefly luciferase CDS. The mRNA fusions were in turn synthesized by *in vitro* transcription from a corresponding linearized plasmid DNA construct using T7 RNA polymerase (Thermo Fisher Scientific), following the manufacturer's instructions. The plasmid DNA was removed after *in vitro* transcription by addition of RNase-free DNase I (Thermo Fisher Scientific). The RNA was purified by phenol–chloroform extraction and dissolved in nuclease free water (Ambion). RNA integrity was assessed on a 1% agarose gel.

For the luciferase assays, the above RNAs were translated in wheat germ extract (Promega), according to the manufacturer's instructions. Each 10 µL reaction consisted of 400–500 ng of RNA and wheat germ extract and was supplemented with 40 µM amino acid mix, 72 mM potassium acetate, and an RNase inhibitor (Thermo Fisher Scientific). Reactions were assembled on ice, followed by pre-incubation at 17 °C for 5 min before shifting to the assay temperatures (17 °C and 27 °C) for 15 min. Reactions were stopped by adding 40 µL of stopping solution (1× PBS, 0.1 µM cycloheximide, 1× protease inhibitor cocktail). 40 µL of the resulting mix was then added to 50 µL of luciferin solution (2 mM D-luciferin, 0.1 mM ATP), and luciferase activity was monitored using a Glomax microplate reader (Promega). The results were analysed on MS Excel. The blank corrected luminescence counts were normalized to corresponding RNA levels and plotted using GraphPad Prism.

#### Supplementary figures

**Figure S1**

**a**, *N. benthamiana* leaves inspected under a UV lamp (UVP Blak-Ray B-100, Fisher Scientific) 5 DPI with *Agrobacterium* carrying an empty vector pHRE or a fluorescence reporter gene construct.

**b**, Fluorometry measurement of the clarified soluble fraction produced from three discs extracted from each area of the leaves in figure 1d. Red (585-15/630-20 nm) and green (470-15/515-20 nm) fluorescence signal is coloured red and green, respectively. The colour of the labels corresponds to the fluorescence gene encoded downstream of the specified 5′ UTR.

**c**, Secondary structure of 75 nt of the 5′ UTR and 75 nt of the reporter CDS predicted on *ViennaRNA RNAfold* [3] at 17 °C and visualized using *forna* [4]. The ThermoSwitch hairpin sequence is coloured blue and the *GFP* or *mLumin* CDS are coloured green or red, respectively.

#### References


[1] H. Peyret, J. K. M. Brown and G. P. Lomonossoff, "Improving plant transient expression through the rational design of synthetic 5′ and 3′ untranslated regions," *Plant methods,* vol. 15, p. 1–13, 2019.

[2] J. Chu, Z. Zhang, Y. Zheng, J. Yang, L. Qin, J. Lu, Z.-L. Huang, S. Zeng and Q. Luo, "A novel far-red bimolecular fluorescence complementation system that allows for efficient visualization of protein interactions under physiological conditions," *Biosensors and Bioelectronics,* vol. 25, no. 1, p. 236–239, 2009.

[3] L. Ronny, S. H. Bernhart, C. Höner zu Siederdissen, H. Tafer, C. Flamm, P. F. Stadler and I. L. Hofacker, "ViennaRNA Package 2.0," *Algorithms for molecular biology,* vol. 6, p. 1–14, 2011.

[4] P. Kerpedjiev, S. Hammer and I. L. Hofacker, "Forna (force-directed RNA): Simple and effective online RNA secondary structure diagrams," *Bioinformatics,* vol. 31, no. 20, p. 3377–3379, 22 June 2015.

### Supplementary figure

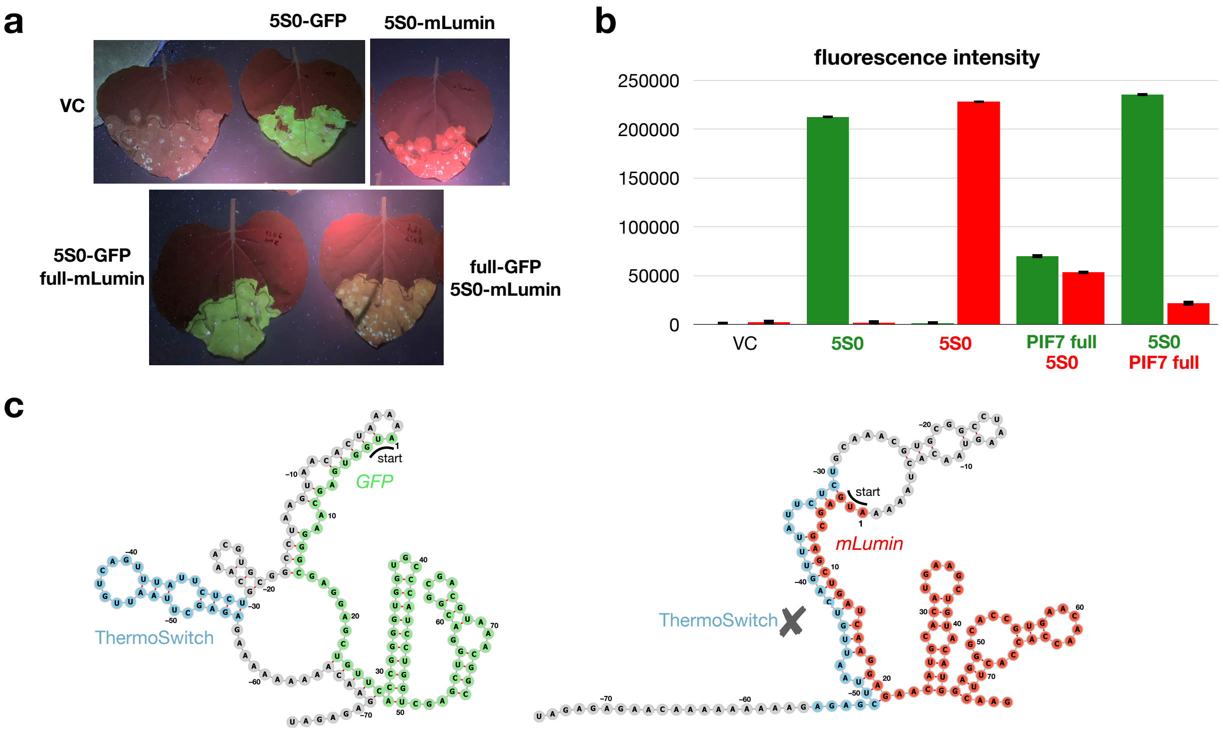

Table S4. 5′ UTR sequences

| 5′ UTR | 5′-UTR sequence |
|---|---|
| **5S0** | TTTAAGAGACGCAACCACAACGCTCTAACGCAATCAATCTACATTATATTAAACGTCTCTAAAA |
| **PIF7 full** | TTTACTGTGATTGTCAAGAGTTTGAACACACAAAGAAGAAAGAAGAACTCAACATTTCAAGCAAGAAGAAAGAGAGAAGAGAGAAGGTCCAA TAATAGAGAGAACAAAAAAAAAGAGAGCTTAATTGTCAGTTTATTCTCTGCAAACGTGCGGCCTAAGTAACACTAAAA |

| primer | primer sequence |
|---|---|
| **35e_SacI_fw** | AGGGAGCTCTGATCACTGAGCATCG |
| **NOS-term_SacI_rv** | AAAGAGCTCAATCGCAACAGCTATGACCATGATTACGC |
